# Freedom of speech predicts animal welfare protection across the globe

**DOI:** 10.1101/2024.11.01.621535

**Authors:** Camille M. Montalcini, Thomas W. Crowther

**Affiliations:** Swiss Federal Institute for Forest, Snow and Landscape Research, 8903 Birmensdorf, Switzerland; Department of Environmental Systems Science, Institute of Integrative Biology; ETH Zürich (Swiss Federal Institute of Technology); Zürich, 8092; Switzerland

## Abstract

The welfare of captive and wild animals is closely tied to human health, nutrition, and environmental wellbeing. Yet, until now, the global determinants of animal welfare remain unclear, as we lack quantitative global data on animal treatment. Here, by collapsing 13 indicators of animal welfare legislation across 50 major animal product-producing countries, we identify two key dimensions reflecting ‘animal welfare protection’ and ‘responsible consumption and production’. While high-income countries with lower dependence on agriculture may have more opportunity to focus on animal welfare, our analysis suggests that the level social freedoms of citizens can dramatically overwhelms this economic signal. We find that the ‘voice and accountability’ of citizens ultimately determines the extent of animal welfare protection, explaining 58% of the variation across the globe. Irrespective of environmental or economic conditions, structures that enhance political freedoms, civil liberties, and citizen participation in governance may provide the necessary foundation for developing legislation to promote animal welfare protection.

## Main text

Most mammals and birds exist within captive settings^1^, while the remainder live in natural environments that are subject to growing human pressure^2^. Directly or indirectly, human decisions influence the welfare of all animals on Earth. Beyond our moral obligation to limit animal suffering, a growing body of evidence suggests that promoting animal welfare can enhance human and environmental wellbeing^3,4^. As such, understanding the factors that determine the quality of animal treatment across the globe is critical if we are going to design strategies to improve the wellbeing of both animals and humans.

In recent decades, theoretical work suggests that the welfare of animals is likely to correlate with various aspects of human and economic development. Specifically, it is expected that economic security may be critical for establishing the financial and political foundations to improve animal treatment^5–8^. With greater access to financial resources, high-income countries are expected to be in a better position to overcome any trade-offs between productivity and animal welfare^6^. Furthermore, in countries that are more dependent on agriculture, the financial risks associated with strict regulation are likely to discourage emphasis on animal welfare policies, whereas countries with lower agricultural dependence may have more opportunity to focus on food quality and animal welfare^5^. However, a growing body of recent research also suggests that cultural and social perspectives may interact with these economic drivers^9–13^, as consumer attitudes may play an increasingly prominent role in regions where people have access to sufficient income^11,14^ and education^14–16^. Indeed, there are some indications that activism to promote animal welfare may have grown in high-income countries, while the need to satisfy basic survival needs is still expected to take priority in low-income regions^6^. However, until now, testing these interacting theories has been precluded by the lack of quantitative global data on animal welfare and its socioeconomic drivers across the globe^5,17^.

Given the highly complex and multidimensional nature of animal welfare, collecting standardised global data on the welfare of captive and wild species in relation to living conditions is practically prohibitive^18^. Therefore, we use a top-down approach to generate standardized animal welfare indicators by compiling data related to policy and legislation commitments across the 50 dominant animal production countries listed by the United Nations Food and Agricultural Organization. By combining 13 indicators representing the intensity of national policy and legislation, along with the level of ‘cruelty’ in meat consumption and production, we could examine the dimensionality of animal welfare commitments of countries. These emergent dimensions represent unified indicators of national-scale animal welfare commitments, which can be linked to various social, economic and environmental variables to examine the determinants of animal welfare across the globe.

Based on our categorical principal component analysis, the scree plots revealed that only three dimensions are necessary to explain 79% of the variation in animal welfare commitments across these 50 countries. Notably, one major dimension was able to explain over 57% of the variation, broadly correlating with all indices relating to the treatment of captive and wild animals (Figure 1b-c). The striking alignment among these variables means that this main dimension can provide a standardized metric to reflect the overall level of policy commitment towards ‘animal welfare protection’. The second orthogonal axis of variation explained 12% of the variation across the dataset, and corresponded with the level ‘responsible production and consumption’ accounting for the number of people in those respective countries and the quality of animal protection laws (Figure 1b-c). While the third axis of dimensionality helped to explain a non-negligible proportion of the variation across countries, it did not strongly align with any specific animal welfare indicators, as all loadings were below 0.5.

**Figure 1.**
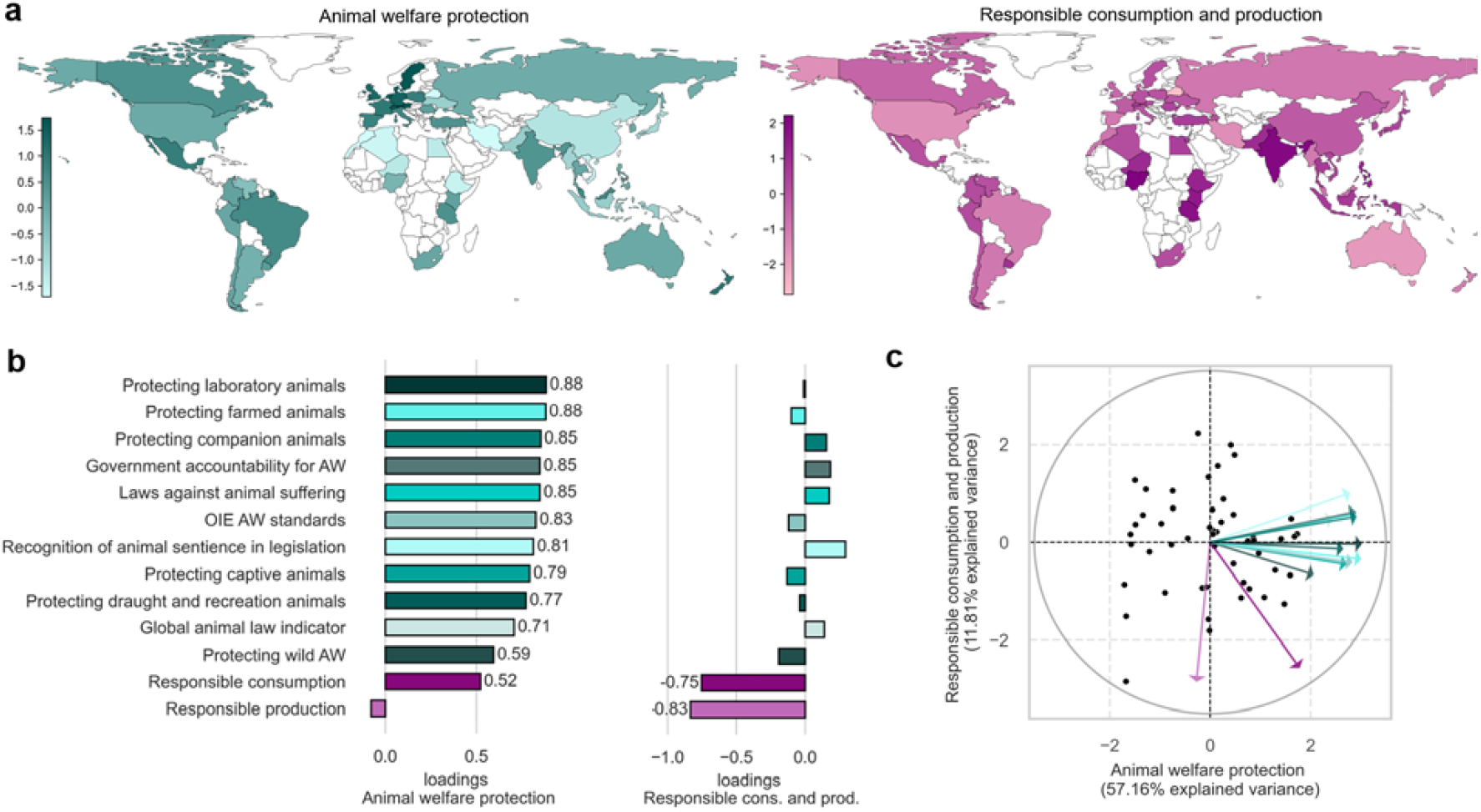
Main dimensions of animal welfare. **(a)** Spatial variation of the two main animal welfare dimensions. **(b)** Loadings of each indicator on the main dimensions, with values shown only for significant loadings (absolute value > 0.5) and ‘AW’ denoting ‘animal welfare’. We ordered the indicators in descending order according to their loadings on the first dimension. **(c)** Biplot of the categorical principal component analysis, where dots represent the raw data.

We projected the data for each country onto the subspace spanned by the two main dimensions to give a score for each of the top 50 animal producing countries (Figure 1a). In terms of overall animal welfare protection (first dimension), the strictest levels of policy commitment were apparent in Europe, followed by New Zealand, and Mexico. However, the second axis – which relates to the per capita level of responsible consumption and production of farm animals – revealed the highest values in India, followed by countries in Central and North Africa. While several of the more developed countries may have stringent animal welfare legislation, they do not appear to demonstrate the most responsible levels of per capita consumption and production of animal products, relative to lower-income countries with lower individual-level meat consumption rates^19^.

We used random forest models to explore how these two main dimensions of animal welfare relate to national-level data on environmental, social and economic conditions. For the main dimension of ‘animal welfare protection’ there was almost no environmental signal, as socioeconomic factors ultimately determined the global pattern (see Figure 2a). However, in contrast to expectations from previous studies^5–8^, the economic indicators such as GDP only explained a relatively small proportion of the variation in animal welfare protection across the globe. While wealthier countries generally tended to have more stringent legislation, there was a large variation within the lower-income regions that could not be explained by economic data (see shaded area of Figure 3). Indeed, there were many countries with relatively low per capita GDP, including Tanzania, India, Kenya, Romania and Brazil, which scored more highly than wealthier counterparts in their commitments to animal protection.

**Figure 2.**
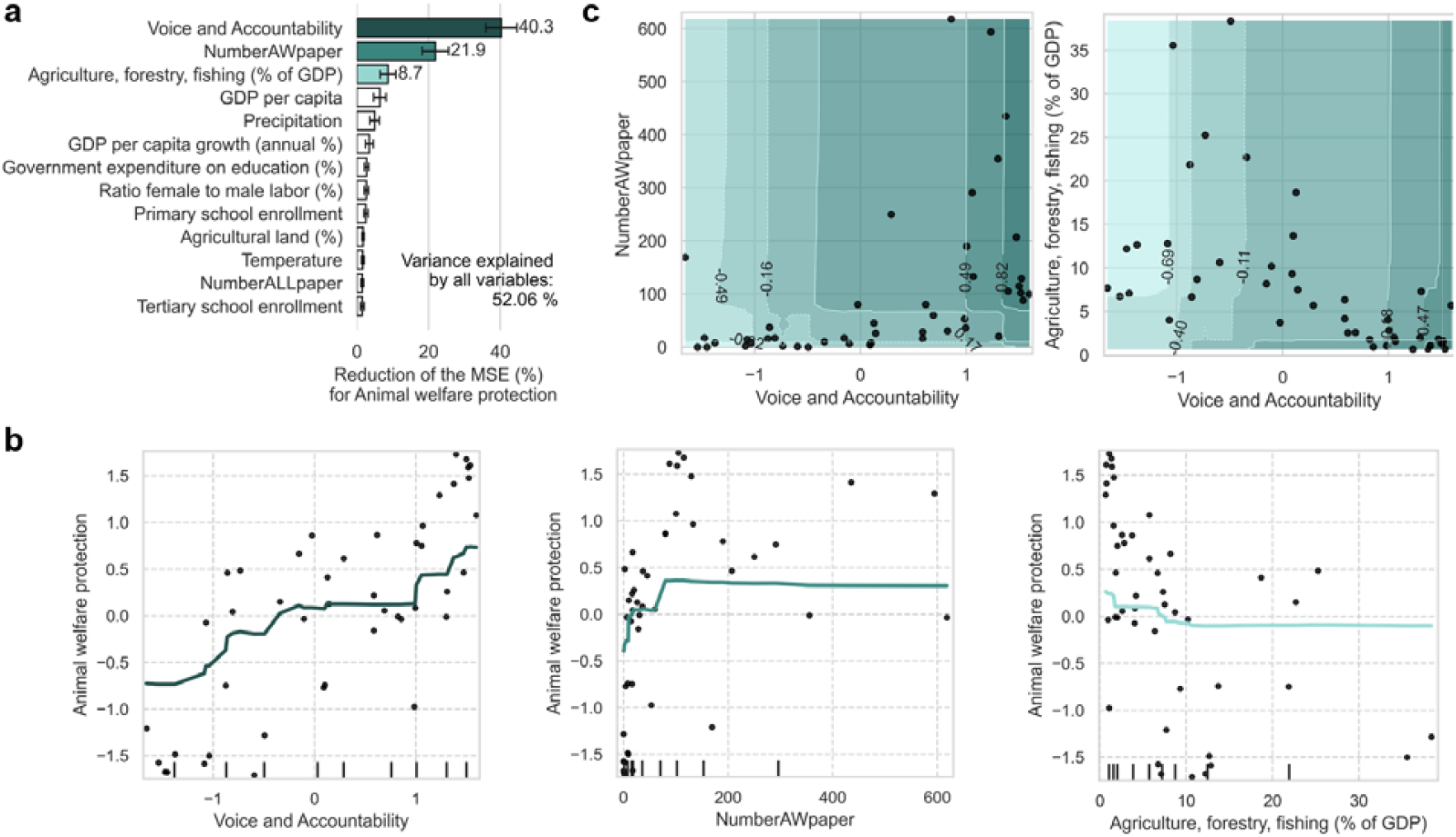
Variable importance and partial dependence plots of the three most important socioeconomic and environmental variables for the first dimension - ‘animal welfare protection’. **(a)** Normalized variable importances from the random forest for animal welfare protection, computed as the total reduction of the mean squared error (MSE) contributed by each variable, so that higher values indicate more important variables. Error bars represent standard deviation. **(b)** Solid lines represent the partial dependence plots of the three most important variables, while black dots indicate the raw data; tick marks on the x-axis illustrate the distribution of each variable. **(c)** Two-way partial dependence plots illustrating potential interactions between the top variable and the next two most important variables, where darker colours would indicate higher animal welfare protection scores. To mitigate extrapolation risk during interpretation we overlaid the raw data on top of the plot.

**Figure 3.**
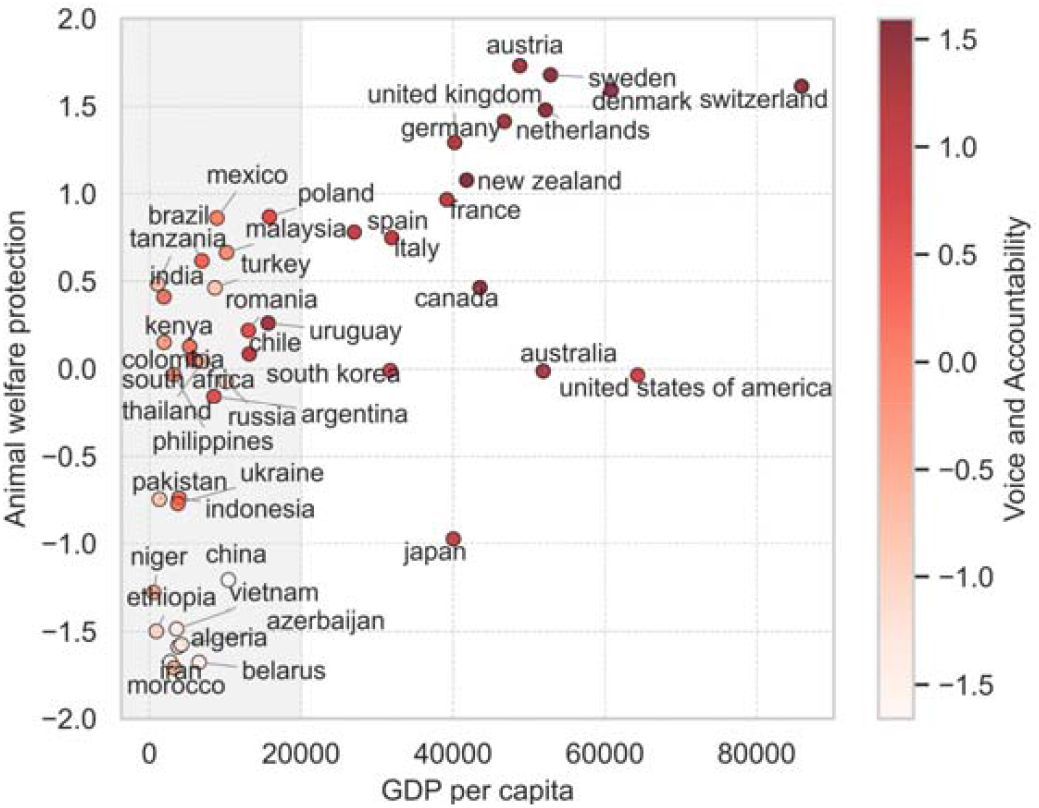
Animal welfare protection in relation to the GDP per capita for all countries used in the analysis, with colours indicating levels of voice and accountability.

By far the strongest predictor of animal welfare protection was ‘voice and accountability’, which broadly reflects the freedom of speech and citizens influence on national policy. This variable dramatically enhanced the performance of the random forests model, reducing the mean squared error by more than twice as much as any other variable. Remarkably, within linear models, voice and accountability captured 58% of the variance on its own (Figure 4), explaining the patterns in animal welfare protection observed across both low- and high-income countries (Figure 3). Specifically, the countries with high voice and accountability consistently showed the greatest tendency to prioritize animal welfare protection, irrespective of economic status. In addition, the number of academic publications related to animal welfare also explained a marginal amount of the variation in animal welfare protection (Figure 2 and 4). This result may reflect a feedback between knowledge and action, as increased scientific knowledge can enhance awareness about animal welfare issues, while greater awareness may also stimulate scientific research to guide public policy^10,13^. However, the potential interactions between the effects of these two main variables (Figure 2c) suggest that academic publishing in animal welfare only serves to amplify the effects of voice and accountability, which dominates the overall signal.

**Figure 4.**
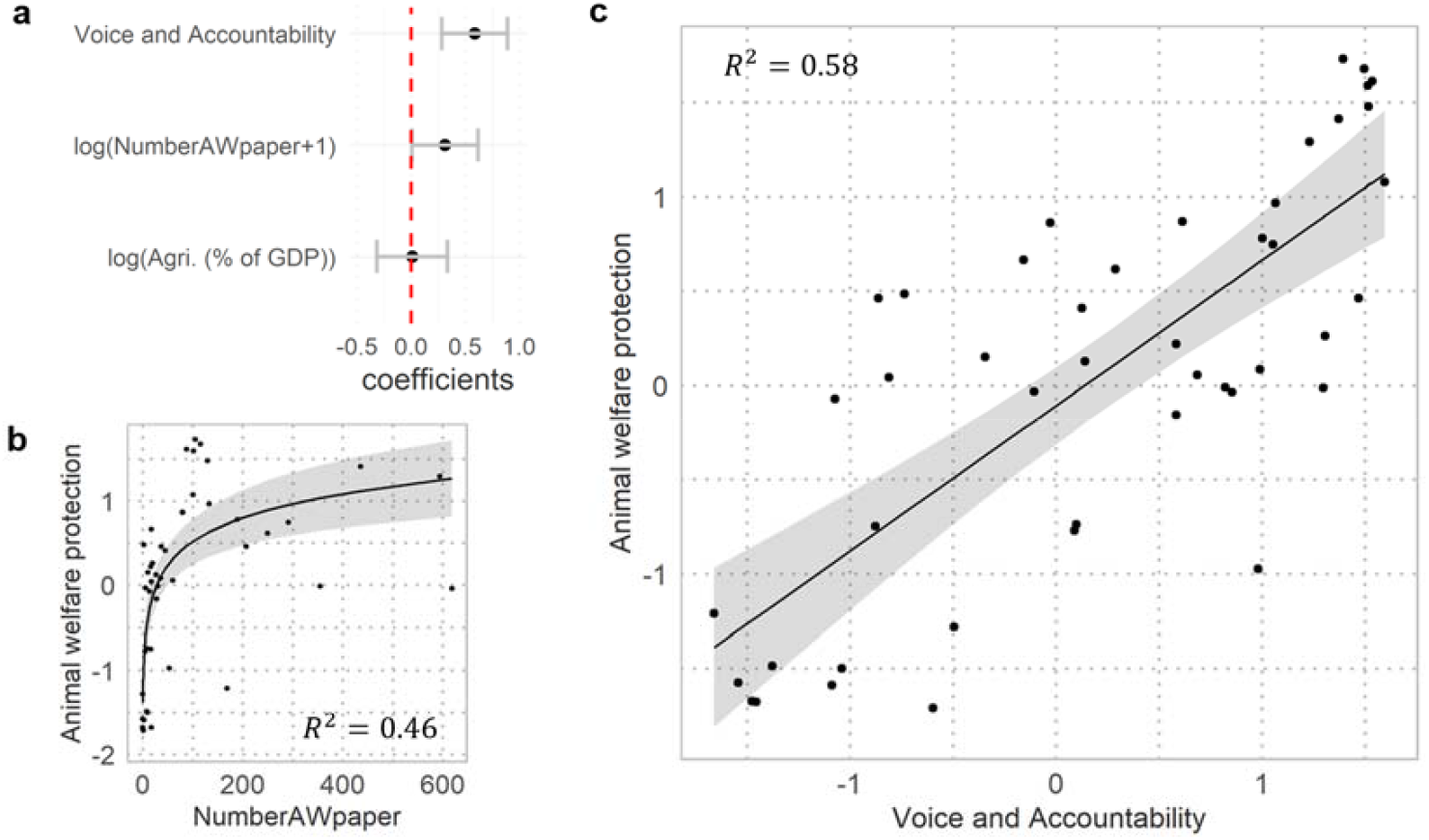
Effect sizes and adjusted predictions for the animal welfare protection dimension derived from the linear regression. **a** Regression coefficients reflecting the change in animal welfare protection for each one standard deviation increase in the predictor variables, and bars indicate the 95% confidence intervals (Voice and accountability: p < 0.001, t value = 0.06; number of publications related to animal welfare (NumberAWpaper): p < 0.05, t value = 2.07). The model included as predictors the three most important variables identified by the random forests, which explained a total of 61.45% of the variance in the animal welfare protection dimension. **b-c** Lines represent the predictions from single-predictor linear regression models using the key variables, the shaded areas indicate the 95% confidence intervals, and the dots represent the raw data.

In contrast to animal welfare protection, we did not identify any strong predictors for the ‘responsible consumption and production’ dimension (Figure 5a). Scores for this second axis generally correlated negatively with tertiary school enrolment, and positively with the economic dependence on agriculture (Figure 5). However, even when accounting for all interactions among environmental and socioeconomic variables, our random forest model could only explain a marginal (approximately 6%) proportion of the variation across the globe. Altogether, these results potentially lends support to prior research, which suggests that changing the behaviour of individual consumers and producers may be more complex than shifting broader national-scale opinion and legislation^13,20^.

**Figure 5.**
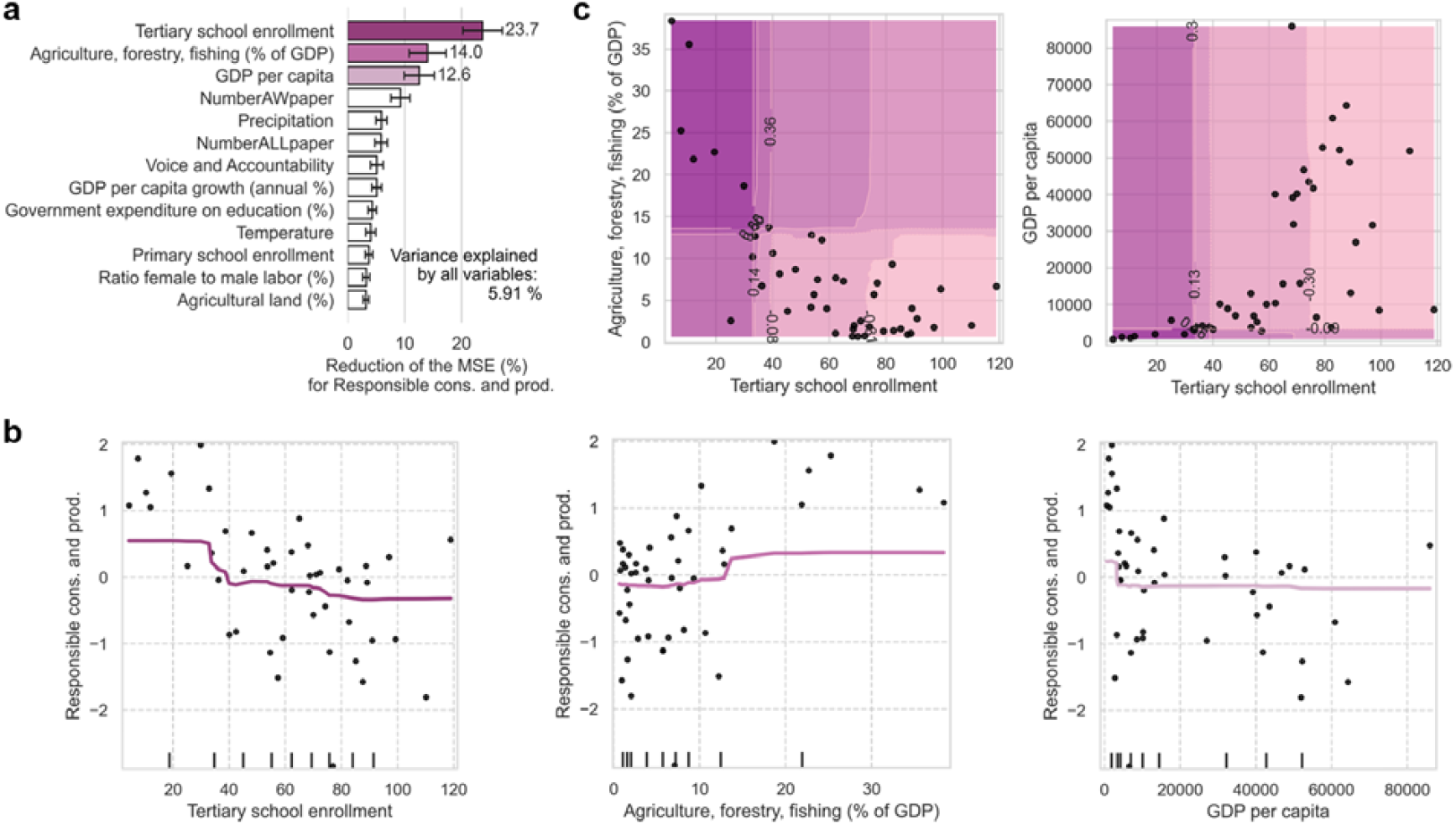
Variable importance and partial dependence plots of the three most important socioeconomic and environmental variables for the second dimension - ‘responsible animal consumption and production’. Similar to Figure 2.

Until now, the highly complex and species-specific nature of animal welfare has precluded efforts to generate standardized data at a global scale^18^. By combining distinct indicators of national policy and legislation, our analysis provides a standardised assessment of policy commitments to animal welfare, revealing two broad axes of variation that relate to ‘animal welfare protection’ and ‘responsible production and consumption’. Our results contradict the contemporary expectation that economic development determines the level of national commitments to animal welfare^5–8^. While high-income countries with lower dependence on agriculture might have a greater freedom to prioritise quality over quantity of animal products, our analysis suggests that the level of citizens’ social freedoms can dramatically overwhelm this economic signal. That is, animal welfare protection is highest in countries where citizens’ opinions are more likely to be reflected in national policy, irrespective of the economic status. This trend potentially suggests that the human condition may generally be favourable towards animal welfare, but that this perspective is only able to emerge in regions where citizens have a greater influence on national policy. Nevertheless, this does not necessarily translate to improved responsibility of consumption and production, which are often highest in lower-income countries.

The correlational nature of our analysis limits our capacity to discern causality in these relationships. It is possible that citizens in countries with high voice and accountability have been responsible for driving strong commitments to animal welfare. However, it is also possible that the countries with higher concern for animal welfare are predisposed to promote societal freedoms of citizens. Future research should examine how these animal welfare indicators change over time in order to establish causality and facilitate effective policy decision making. Nevertheless, the tight coupling of animal welfare protection with social empowerment lends support to the idea that there may be common avenues for tackling animal welfare conditions in both low- and high-income countries^6^. Irrespective of the economic context, social structures that enhance political freedoms, civil liberties, and citizen participation in governance may lay the necessary foundations for developing legislation to promote the protection of animals across the globe. Ultimately, this highlights potentially bright policy avenues for simultaneously improving animal and human wellbeing at national and international scales.

## Methods

### Main dimension of animal welfare

We used 13 indicators of animal welfare derived from three data sources, covering 50 countries selected from the largest producers of farmed animal products in the world. First, we used 10 animal protection indicators ranking the countries according to their legislation and policy commitments to protecting animals, published in 2020 by the ‘World Animal Protection International’ global organisation^21^. The 10 indicators cover four main aspects: the recognition of animal sentience and prohibition of animal suffering (two indicators), the establishment of supportive government bodies (one indicator), and the support for international animal welfare standards (one indicator), and the presence of animal welfare legislation (six indicators: farm animals, companion animals, animals used for draught and recreation, laboratory animals, wild animals, and captive animals which includes zoo animals, private keeping of wild animals, and fur farming). The indicators scores countries from A (being the highest score) to G (being the weakest score). Secondly, we used the national database on animal legislations from the ‘Global Animal law Association’, based on the available online information as of 31st of March 2021^22^. Lastly, we used two indicators from the ‘voiceless animal cruelty index’ (VACI) developed by the ‘Animal Protection Institute’ in 2017 and updated in 2020^23^. These two indicators assessed countries based on the nature, extent, and severity of farmed animal cruelty while considering the quality of animal protection laws. Specifically, the ‘producing cruelty’ indicator, hereafter referred to as ‘responsible production’, scored countries based on the number of farmed animals slaughtered per capita, accounting for variations in animal treatment and protection. The ‘consuming cruelty’ indicator, hereafter referred to as ‘responsible consumption’, evaluates countries based on their consumption of farmed animals, examining the number of animals consumed per capita and the ratio of plant-based protein to farm-animal protein consumed.

Since we have both ordinal and continuous indicators, we performed a categorical principal components analysis (CATPCA)^24^ to identify the main dimensions explaining most of the variance. This method transforms the original ordinal variables into a numerical scale that maximize the variance explained by each component^25^. Therefore, the number of dimensions affects the results, making it important to compare results across different dimensionalities when choosing the number of components. Thus, using the ‘gifi’ package and its ‘princals’ function^26^, we examined scree plots for solutions ranging from two to six dimensions, suggesting three components. Before performing any analysis we discretized the responsible production and consumption indicators by standardizing them, multiplying all values by 10, rounding, and adding a constant so that the lowest resulting integer value equals one^24^. We then projected all observations onto the subspace spanned by the identified components to obtain one component score for each country per dimension.

### Drivers of the main animal welfare dimensions

To identify the socioeconomic factors contributing most to the identified animal welfare dimensions, we selected a total of 16 variables to capture key factors that may relate to various aspects of animal welfare, as suggested by prior literature. First, we selected 12 variables from the world development indicators^27^ including variables related to agriculture, education, economics, governance, and gender equality. Women tend to have a more positive and compassionate attitude towards animals compared to men^28–30^, so that countries with lower gender inequality could have stronger animal protection policies. Furthermore, as scientific research on animal welfare may guide public policy to protect animal welfare^10,13^, we included the number of publications related to animal welfare (‘NumberAWpaper’) and controlled for the total number of scientific publications. Lastly, we included environmental factors (average mean surface air temperature and precipitation from the Climate Change Knowledge Portal^27^) as they can influence political decisions and exert additional abiotic pressures on production^8^.

As education variables, we selected the percentage of government expenditure used on education and the school gross enrolment ratio for primary, secondary, and tertiary education, defined as the ratio of total enrolment to the population of the relevant age group. As economic variables, we selected the gross domestic product (GDP) per capita, representing countries GDP divided by their total population, and the annual percentage growth rate of GDP per capita. As agricultural variables we selected the percentage of land area that is used as agricultural land, and the value added of the agriculture, forestry, and fishing to GDP (in percentage). As governance variables we selected the ‘Voice and Accountability’ capturing ‘*perceptions of the extent to which a country’s citizens are able to participate in selecting their government, as well as freedom of expression, freedom of association, and a free media*’, the ‘Rule of Law’ capturing ‘*perceptions of the extent to which agents have confidence in and abide by the rules of society, and in particular the quality of contract enforcement, property rights, the police, and the courts, as well as the likelihood of crime and violence*’, and the ‘Government Effectiveness’ capturing ‘*perceptions of the quality of public services, the quality of the civil service and the degree of its independence from political pressures, the quality of policy formulation and implementation, and the credibility of the government’s commitment to such policie*’^31^. Finally, we selected the ratio of female to male labour force participation rate (in percentage) as gender equality variable, where labour force participation rate refers to the proportion of the population ages older than 14 that is economically active.

To account for scientific research on animal welfare, we used the total number of articles and reviews published from 2020 to 2021 (to account for the delay associated with the publication process) indexed in Web of Science that included ‘animal welfare’ or related synonyms (i.e., animal wellbeing, animal well-being, animal affective state, animal emotion, animal mood, and animal affect) in their abstracts, titles, or keywords. We used only records with at least one country or region as provided by Web of Science, resulting in 3,466 records. To control for general scientific knowledge, we similarly extracted the total number of articles and reviews per country by iteratively adding the most common written English words (as given in ^32^, p.181) as keywords in a full-record search until the addition of a new word contributed fewer than 1,000 new records. This process resulted in a total of 5,330,718 records, using *the, of, and, a*, and *in* as keywords.

As first preprocessing step, we replaced missing values for the year 2020 for any data from the world development indicators with the mean values from 2019 and 2021. When these values were unavailable, we used linear interpolation based on the two most recent preceding years, excluding any data prior to 2017. We removed variables with more than 10% of missing data, namely, school enrolment for the secondary. To avoid multicollinearity, we iteratively removed the variable with the highest variance inflation score and recomputed the scores until all were strictly below five, using the ‘Statsmodels’ python library^33^. This process led to the removal of the ‘Government Effectiveness’ and ‘Rule of Law’ indicators (see Text S1 for additional analysis including these two variables). Lastly, we excluded any countries with at least one missing value, specifically, Myanmar, Egypt, Nigeria, Peru, and Venezuela. Data used for subsequent analysis included 45 countries and 13 variables, with only three having imputed data.

To contrast the impacts of these socioeconomic and environmental variables on the identified animal welfare dimensions, we used random forests, as they work well with small sample size that have a relatively large number of variables and non-linear data structures^34^. We used the Breiman’s Random Forest method^35^ as implemented in the Scikit-learn python library^36^. Model performance was assessed using leave-One-Out Cross-Validation a procedure well adapted to small dataset^37^ where each observation is considered once as the testing set, with the others as the training set. We evaluated model performance using the explained variance and used 10 different random seeds. For each model fit, we extracted the normalized feature importances computed as the total reduction of the mean squared error brought by each variable, so that higher values indicate more important variables, and reported the mean ± standard deviation across all fits. To visualize the relationships between each variable and the animal welfare dimensions we used partial dependence plot. These plots show the dependence between the identified animal welfare dimensions and one or two variables of interest, marginalizing over the values of all other input variables. To evaluate the robustness of the results across different statistical approaches, we fitted a linear regression model with the key variables identified by the random forest and compared their effect sizes. Finally, we evaluated the variance explained by each key variable on their own, using single-predictor linear regression models. We verified model assumptions, including normality and homoscedasticity of residuals, and log-transformed variables when needed to ensure linearity in their relationships with the animal welfare dimension.

## Supporting information

Test S1

## Data and code availability

All datasets used in this paper are available online, as referenced in the methods. The identified animal welfare dimensions and code are provided in the supplementary information (see Zenodo repository upon publication).

## Author contributions

C.M.M and T.W.C contributed the conceptualization of the project, the methodology, investigation and project administration. C.M.M contributed to the visualization. T.W.C. obtained funding.

