## Supplementary material for "Freedom of speech predicts animal welfare protection across the globe": Test S1

### Text S1

Since *rule of law* and *government effectiveness* were removed during the variable selection procedure due to high VIF, and both are correlated with our strongest predictor *voice and accountability*, we also examined their importance in explaining the overall animal welfare protection. We did this by considering each variable once individually and once in combination with the other predictors. Specifically, we first fitted one random forest model with the selected variables, replacing *voice and accountability* with either *rule of law* or *government effectiveness*. Then, we fitted one separate random forest model for each of these three variables alone. We ran each model with 10 random seeds and report the mean and range (minimum and maximum) of the variance explained by the set of predictors in Table S1. These results confirm that *voice and accountability* is the most important predictor of our animal welfare protection dimension. Interestingly, they also show that a model based solely on this predictor outperform the one that included all socioeconomic and environmental variables, suggesting that models with fewer predictors should be considered for prediction purposes.

| **predictors** | **Single predictor** | **Multi-predictor** |
|---|---|---|
| Government effectiveness | 0.09 (0.07, 0.11) | 0.43 (0.41, 0.45) |
| Rule of Law | 0.24 (0.21, 0.27) | 0.46 (0.44, 0.48) |
| Voice and Accountability | 0.60 (0.59, 0.60) | 0.52 (0.50, 0.54) |

**Table S1** – Mean and range (minimum, maximum) of variance explained by random forest models using different set of predictors.
